# Size- and sex-specific predation on dung flies by amphibian and arthropod predators – size match matters

**DOI:** 10.1101/631549

**Authors:** Wolf U. Blanckenhorn, Gabriele Cozzi, Gregory Jäggli, Juan Pablo Busso

## Abstract

Because predator-prey interactions in nature are multifarious, linking phenomenological predation rates to the underlying behavioural or ecological mechanisms is challenging. Size- and sex-specific predation has been implicated as a major selective force keeping animals small, affecting the evolution of body size and sexual size dimorphism. We experimentally assessed predation by various amphibian (frogs and toads) and arthropod predators (bugs, flies, spiders) on three species of dung flies with similar ecology but contrasting body sizes, sexual size dimorphism and coloration. Predators were offered a size range of flies in single- or mixed-sex groups. As expected based on optimal foraging theory, some anurans (e.g. *Bufo bufo*) selected larger prey, thus selecting against large body size of the flies, while others (*Bombina variagata* and *Rana esculenta*) showed no such pattern. Small juvenile *Rana temporaria* metamorphs, in contrast, preferred small flies, as did all arthropod predators, a pattern that can be explained by larger prey being better at escaping. The more mobile males were not eaten more frequently or faster than the cryptic females, even when conspicuously colored. Predation rates on flies in mixed groups permitting mating activity were not higher, contrary to expectation, nor was predation generally sex-specific. We conclude that the size-selectivity of predators, and hence the viability selection pattern exerted on their prey, depends foremost on the relative body sizes of the two in a continuous fashion. Sex-specific predation by single predators appears to contribute little to sexual dimorphism. Therefore, the mechanistic study of predation requires integration of both the predator’s and the prey’s perspectives, and phenomenological field studies of predation remain indispensable.

## INTRODUCTION

Predation is a fundamental ecological process that plays a central role in the evolution of animal life histories and affects most if not all species (Roff, 1992). As such predation features prominently in life history optimality (e.g. Stearns & Koella, 1986; Fraser & Gilliam, 1992; Werner & Anholt, 1993; Abrams *et al.*, 1996; Leimar, 1996; Kozlowski, 1992; Ruxton *et al.*, 2004) as well as food web and functional response theory (Kalinkat *et al.*, 2011, 2013). Though ubiquitous, predation is not easily assessed beyond the traditional and very crude food web perspective (Cohen *et al.*, 1993; Brose, 2010). The major problem is that most predators, contrary perhaps to many parasitoids (e.g. Stireman & Singer, 2003; Sandrock *et al.*, 2011), are typically not very specialized on particular prey species (Symondson *et al.*, 2002; Ruxton *et al.*, 2004; Scherer & Smee, 2016). Therefore predator-prey interactions in nature are necessarily complex, depending on habitat, ecological context, body size or taxonomic group, to only name the most obvious factors. As a consequence, it is difficult to link phenomenological predation rates to the underlying behavioural or ecological mechanisms, as called for by Blanckenhorn (2000, 2005) when reviewing experimental studies of viability selection. This in turn is problematic for generating realistic estimates for predictive ecological models in theoretical and applied contexts (Kalinkat *et al.*, 2011; Remmel *et al.*, 2011).

Size- and sex-specific predation has been suggested as a major selective force keeping animals small and thus affecting the evolution of body size and sexual size dimorphism (Blanckenhorn, 2000, 2005). Whereas the evidence for sexual and fecundity selection favouring larger body size in males and females is overwhelming, counterbalancing selection forces disfavouring large body size remain cryptic. If a majority of predators had reported preferences for larger prey this problem might be solved. Importantly in this context, mortality risk of most animals is mediated by behaviour at any life stage in many contexts: the hunting behaviour of the predator, the evasion behaviour of the prey, or the foraging behaviour of either necessary to avoid starvation (Ruxton *et al.*, 2004; Blanckenhorn, 2005). Optimal foraging theory postulates and documents that predators are sensitive to the energy content and size of their prey (Stephens & Krebs, 1986; Sih & Christensen, 2001). On the other hand, at one point prey become too big and strong for any given predator requiring excessive handling time, or might no longer fit into the predator’s mouth (the concept of gape limitation: e.g. Truemper & Lauer, 2004), implying an optimal prey size in terms of profitability (i.e. energy obtained relative to energy invested: Stephens & Krebs, 1986; e.g. Elner & Hughes, 1978). Therefore, a rough positive correlation between predator and prey size and, globally, intermediate predator-prey size ratios are expected (Brose *et al.*, 2006). Taking the prey’s perspective, however, studies often find viability advantages of larger individuals, rather than the expected disadvantages based on optimal foraging of the predator (Blanckenhorn, 2000, 2005). This relates to some extent to the hypo-allometric scaling of metabolic rate (Reim *et al.*, 2006; Blanckenhorn *et al.*, 2007), but surely also involves gains in strength aiding larger prey in escaping predation. Moreover, as they transition through life stages, growing prey will likely consecutively pass through the optimal size windows of various predators of ever larger size, facing predation risk at most times (Wellborn, 1994; Berger *et al.*, 2006; Mänd *et al.*, 2007; Remmel *et al.*, 2011). It follows that the mechanistic study of predation requires integration of both the predator’s and the prey’s perspectives, but even when doing so might never be complete because the entire spectrum of prey or predators cannot easily be covered.

Experimental studies of sex- and size-selective predation are uncommon but exist in the literature. The topic seems to be of more interest in aquatic (fish, cladocerans, tadpoles, etc.) than terrestrial organisms (e.g. Sogard, 1997; Scherer & Smee, 2016). Terrestrial investigations in the functional response and food web context concentrate on soil invertebrates (Kalinkat *et al.*, 2013) but are typically phenomenological in frequently ignoring crucial behavioural elements (but see Vucic-Pestic *et al.*, 2010; Kalinkat *et al.*, 2011, for good counterexamples). Remmel *et al.* (2011) recently reviewed predation on holometabolous insect larvae feeding on plant leaves (folivory) and found that birds preyed predominantly on larger larvae, while arthropod predators rather preferred smaller prey (see also Sogard’s (1997) review for marine fish). This result emphasizes that when taking the predator’s perspective, we may obtain a clear pattern, but when taking the prey’s perspective net selection on body size may well turn out to be unrelated to size because usually several predators prey on the same species, some of which will prefer small and others large individuals (e.g. Lüning, 1992; Rice *et al.*, 1993).

Apart from body size, the sex of the prey should also matter. Even when of the same size, males and females of a given species can behave very differently, ultimately affecting their probability to be preyed upon. For example, predation risk has been found to be greater for the sex attracting mates via acoustic or visual signals, or for the sex that has to move more or farther to find mates. Thus, the sex competing more for mates, typically males, frequently faces greater predation risk or energy expenditure when searching (Slagsvold *et al.*, 1988; Magnhagen, 1991), attracting (Partridge & Farquhar, 1981; Lima & Dill, 1990; Cordts & Partridge, 1996; Kotiaho *et al.*, 1998; Zuk & Kolluru, 1998), assessing (Pomiankowski, 1987; Hedrick & Dill, 1993), or rejecting potential mates (Rowe, 1994; Watson *et al.*, 1998; Jormalainen *et al.*, 2001). Moreover, the act of mating itself, or that of avoiding mating, should also increase predation rates, if only because a mating pair attracts more attention, is more visible and/or less mobile (Blanckenhorn *et al.*, 2002; Mühlhäuser & Blanckenhorn, 2002). It is therefore likely that sex and body size interact in determining the outcome of predation, for instance if large prey individuals are more likely to mate (which is common), if there is size-assortative mating, or if predators systematically only eat one of the mating partners because of its position (Teuschl *et al.*, 2010).

Pastoral grasslands are ubiquitous habitats harbouring a large array of interacting species that typify the agricultural landscape of many developed countries. We here investigate predation on three of the most abundant and widespread dung flies in Europe (Pont & Meier, 2002; Blanckenhorn *et al.*, 2010) with similar ecology but contrasting body sizes, sexual size dimorphism and coloration (i.e. conspicuousness). Yellow dung fly (*Scatophaga stercoraria*) males are bright yellow to orange, hairy and on average considerably larger than the greenish, more cryptic females. *Sepsis cynipsea* males are smaller than females and both shiny black; *S. thoracica* males feature an additional male dimorphism with large orange and small black males, and black females of intermediate size (Busso & Blanckenhorn, in review). All these flies lay their eggs into cow dung, in which the larvae develop. Their mating behaviour around the dung of livestock (primarily cows) is characterized by scramble and contest competition of males for access to females, with at times conspicuous mate rejection and struggle behaviour, and relatively long copulation and guarding durations (20 – 60 min: Parker, 1970, 1972; Ward, 1983; Ward *et al.*, 1992; Blanckenhorn *et al.*, 2000; Jann *et al.*, 2000). All dung fly species also spend considerable time foraging for nectar and insect prey (*Sc. stercoraria* only) in the areas surrounding pastures (Warncke *et al.*, 1993), forested or not. Accordingly, in addition to various specialized predators of juvenile stages in and around the dung (Hammer, 1941; Skidmore, 1991; Pont & Meier, 2002), adult dung flies regularly face a large guild of generalist predators in their habitat. These include vertebrates such as various insectivorous birds, *Lacerta* lizards, and several amphibians where ponds and streams are common, but also many invertebrates, primarily jumping, crab and orb-weaver spiders and predatory insects (wasps, scorpion flies, bugs, various dipterans including adult *Sc. stercoraria* itself, etc.; Hammer, 1941; pers. obs.). Unfortunately, no systematic assessment of the predator guild is available.

Our aim was to obtain a more complete picture of size- and sex-specific predation in another terrestrial system so as to generalize, verify or refute the patterns found for folivorous insects (Remmel *et al.*, 2011). Our selection of predators and prey is necessarily idiosyncratic and incomplete, however should be representative for the common pastoral habitat considered. We expected that the larger vertebrate predators (mostly anurans) would prefer larger flies when feeding on relatively small insect prey so as to optimize their energy gain, whereas the smaller arthropod predators (bugs, flies, spiders) would rather select smaller individuals (if selective at all) as larger prey require prohibitive handling (Brose et al., 2006). We further expected the more mobile and often more colourful males to be subject to higher predation than the cryptic females, and flies in a mixed setting permitting mating activity to be subject to higher predation than in single-sex groups (Remmel & Tammaru, 2009; Svanbäck & Eklöv, 2011; Teuschl *et al.*, 2010).

## MATERIALS AND METHODS

### Prey species

As prey we used three species of dung flies: the yellow dung fly *Scathophaga stercoraria* (Diptera: Scathophagidae), and the two black scavenger fly species *Sepsis cynipsea* and *S. thoracica* (Diptera: Sepsidae). The sepsids are primarily black and considerably smaller than scathophagids (6 – 13 vs. 3 – 9 mm in length; 8 – 48 vs. 0.5 – 3.6 mg in wet mass). The first two species are the most common dung flies in north-central Europe and have been used extensively as models in evolutionary ecology (see Blanckenhorn, 2007, for exemplary references; Blanckenhorn *et al.*, 2010 and http://sepsidnet-rmbr.nus.edu.sg/ for pictures). *Sepsis thoracica* is the most common sepsid fly around livestock dung in southern Europe, but is also frequent north of the Alps (Pont & Meier, 2002).

Prey flies used in this experiment all stemmed from laboratory cultures held in groups (sepsids) or individually (*Sc. stercoraria*) in plastic or glass containers under standard conditions with water, sugar as food, and cow dung as food and oviposition substrate (see Blanckenhorn et al. 2000, 2010, for details). To obtain a broad range of prey body sizes, the phenotypic size of the flies was manipulated by raising them at various dung (i.e. food) quantities and/or population densities. The hind tibia length of all prey was determined under a binocular microscope and used as an estimate of their body size. All prey individuals were used only once.

### Vertebrate predators

As vertebrate predators we used a number of amphibian species of various sizes common in grasslands in Switzerland. In general, we collected predator individuals ourselves around Irchel campus, a typical Swiss pasture habitat close to forest featuring some ponds and streams (47.40°N, 8.55°E). Some individuals stemmed from experimental populations colleagues held on campus. Snout-vent-length (SVL) of all predators was used as standard estimate of their size, using calipers. Amphibian predators were held in 1.3 × 1.3 × 1.8 dm^3^ plastic containers (cf. below) and used repeatedly. While in the laboratory we fed each predator with a mix of *Drosophila* species (*D. melanogaster, D. virilis, D. americana*, *D. novamexicana*). At the end of our experiments predators were released into the wild where originally captured.

We originally attempted a crossed experimental design in confronting all predators with both *Sc. stercoraria* (large prey) and *Se. cynipsea* (small prey; *Se. thoracica* data were added later, see below). However, most amphibian predator species ate only one or the other prey species to yield a complete data set (Table 1), and the marsh frog *Rana ridibunda*, the Alpine newt *Triturus alpestris*, and the fire salamander *Salamandra salamandra* did not produce usable data sets for either prey species in captivity. Complete data sets with at least ten replicates per sex treatment (explained below) were obtained for the common toad *Bufo bufo* (4 individuals, size range 55 – 82 mm SVL; predation experiments with *Sc. stercoraria* and *Se. cynipsea*), the yellow-bellied toad *Bombina variegata* (11 individuals, size range 26.9 – 37.7 mm SVL; *Sc. stercoraria* only), the edible frog *Rana esculenta* (now *Pelophylax esculentus*; 8 individuals, size range 58 – 74.5 mm SVL; *Sc. stercoraria* only), and juveniles of the common frog *Rana temporaria* (4 individuals, size range 13.6 – 14.7 mm SVL; *Se. cynipsea* only; Table 1). All these amphibians are largely sit-and-wait predators but also actively orient towards moving prey.

**Table 1.**
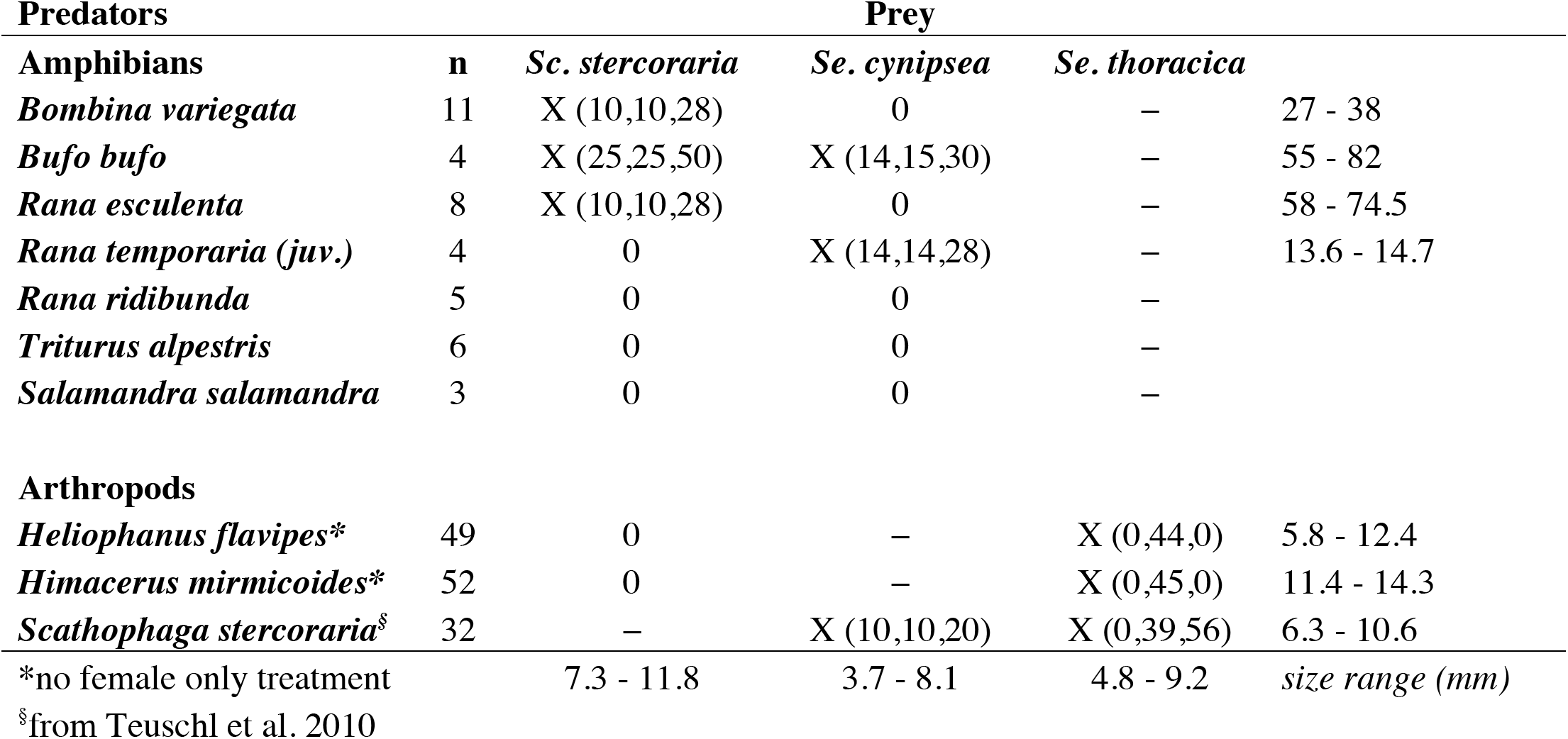
Combinations of predators and prey used (X) with number of tested replicate groups (of females, males, mixed) in parentheses and size ranges (snout vent length for amphibians; total length for arthropods). 0: attempted but too little predation; −: not attempted.

### Arthropod predators

As arthropod predators are small relative to their prey, we only used sepsids as prey for them. (While *Sc. stercoraria* would cannibalize individuals of their own species regularly, the other two predators would not regularly manage to catch this large prey species.) We in the end selected three arthropod predators differing by taxon, in body size and hunting technique, after verifying in the field that they in fact prey on sepsids: the jumping spider *Heliophanus flavipes* (Areanae: Salticidae), an active hunter 5 – 13 mm in total length; the damsel bug *Himacerus mirmicoides* (Heteroptera: Nabidae), a sit-and-wait predator 11 – 15 mm long; and the predatory yellow dung fly *Scathophaga stercoraria* (Diptera: Scathophagidae), a sit-and-wait but also active hunter 6 – 12 mm long. Predation experiments with *Sc. stercoraria* as predator and *Se. cynipsea* as prey were performed around the time of the experiments with amphibian predators described above (cf. Teuschl *et al.*, 2010), whereas those using the arthropod predators and *Se. thoracica* were carried out more recently.

The spiders and bugs were collected around Irchel campus and subsequently maintained individually in 1 × 1 × 1.4 dm^3^ plastic containers (cf. below) in the laboratory. We fed each predator with water and a mix of *Drosophila* species (*D. melanogaster, D. virilis, D. americana*, *D. novamexicana*). Only adult arthropods were used as predators, each individual only once. *Sc. stercoraria* individuals were raised in the laboratory as mentioned above.

### Size-dependent predation experiments

To investigate size- and sex-dependent predation, we set up single-sex and mixed-sex groups of flies of varying body size. We examined 10 – 28 replicate groups per predator and sex treatment. In the experiments with amphibian predators, we randomly assigned 10 males, 10 females, or 5 males plus 5 females (in the mixed-sex case) prey individuals of various sizes (assessed by eye) into a transparent 1.3 × 1.3 × 1.8 dm^3^ experimental container. Because all amphibians swallow their prey whole, all flies were measured before the experiment, and only the surviving flies were measured again thereafter (to match individuals and calculate selection coefficients).

In the experiments with arthropod predators and male *Se. thoracica* as prey, we assembled 3 (large) orange plus 3 (small) black males into smaller 1 × 1 × 1.4 dm^3^ transparent experimental containers. The corresponding mixed-sex treatment consisted of 4 males plus 4 females, whereby 10 replicates featured only large orange males, 10 only small black males, and 10 both orange and black males (2 + 2). No female only treatment was conducted with *Se. thoracica* as prey as we were primarily interested in the effect of coloration at this point. In the experiment with *Sc. stercoraria* preying upon *Se. cynipsea*, 16 prey individuals were used in the above-mentioned larger containers (see Teuschl et al. 2010). All the arthropod predators used suck the content out of their victims (extra-intestinal digestion), leaving the entire exoskeleton behind so it could be measured after the experiment.

All test containers were provided with water, sugar, pollen and fresh cow dung, plus a natural structure (plant leaf, branch, etc.) as potential shelter for the predators and the prey. For the anuran predators we additionally added half a plant pot as shelter, a water dish and some soil. In each single replicate, the prey flies were released into the test containers first and then given at minimum 15 min to accommodate. The predator was added thereafter, marking the beginning of the experimental period. Predators were starved for varying amounts of time (few hours to days) before the experiment so as to be hungry, whereby their hunger level was not of interest here as long as they caught prey. Experimental replicates were stopped when approximately half of the flies were eaten, and the time this took was noted. This temporal calibration of the experiment was necessary because the predators varied strongly in how fast they ate their prey depending on species, hunger level, etc., and because selection coefficients cannot be computed if none or all prey survived.

### Statistical analysis

We used the standard methods of Arnold & Wade (1984) for dichotomous fitness measures (dead/alive = 1/0) to estimate selection differentials. For each prey group all tibia lengths (x) were first *z*-scored as *z*_*i*_ = (*X*_*i*_ – mean(X)) / *SD(X)*, thus standardizing mean and variance. In the mixed sex groups females and males were *z*-scored separately because of size dimorphism (Teuschl et al. 2010). The selection differential is then simply equal to the mean standardized body size of the surviving flies. Note that this standardization works equally for any (linear) trait considered as size measure, and that the total number of prey and their size range within the group, which here varied depending on predator, prey, treatment and container size, only affects the precision of the selection differential but does not bias the estimate, thus permitting direct comparisons of naturally varying group sizes and species (Arnold & Wade, 1984).

The resulting selection differentials were analysed as outcome variables using general linear models with predator and prey species, treatment (females only, males only, mixed sex) and sex within treatment as fixed factors. For the amphibians that were used repeatedly, predator individual was additionally entered as random effect. In such models the intercept can serve to test for consistent selection across treatments, predator and/or prey species (cf. Fig. 1).

**Figure 1.**
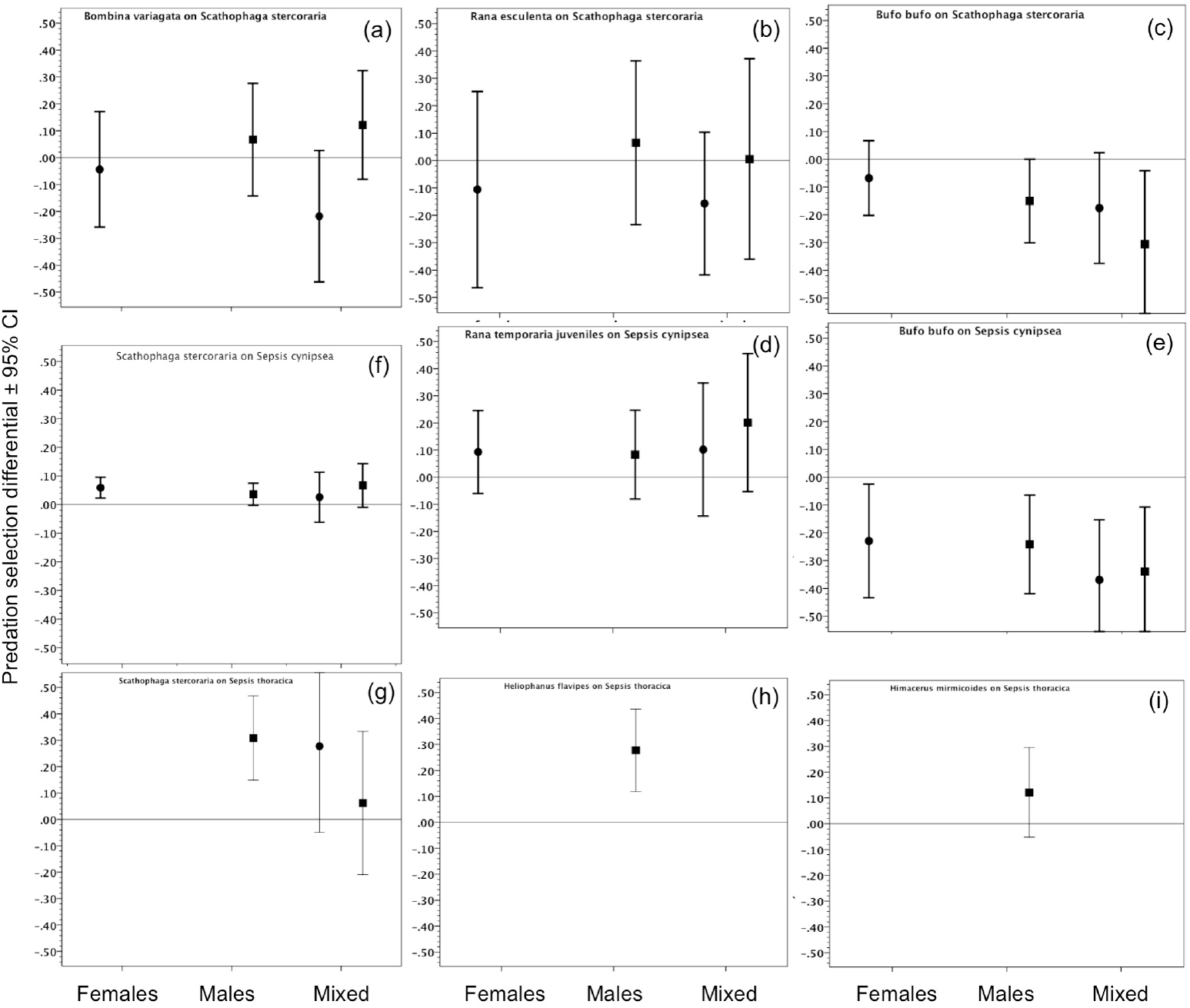
Standardized mean viability (i.e. predation) selection differentials (± 95% CI) as a function of predator ((a-e) amphibian vertebrates; (f-i) arthropods) and treatment (all-male, all-female, mixed-sex). Males are represented as squares and females as circles.

## RESULTS

Amphibian predators showed no systematic size selectivity regarding fly prey, whereas the three arthropod predators generally ate more small flies, thus overall exerting positive viability selection (Fig. 1). The 95% confidence intervals of several individual selection differentials overlap zero in Fig. 1 indicating no significance. Overall significant negative selection against large prey was exerted only by the toad *Bufo bufo* on both *Sc. stercoraria* and *Se. cynipsea* (significant intercepts in the separate models including all treatments (females, males, mixed): *F*_1,96_ = 10.20, *p* = 0.002 and *F*_1,54_ = 24.52, *p* < 0.001, respectively; Fig. 1c,e). Overall significant positive selection against small prey flies was exerted by *Rana temporaria* juveniles on *Se. cynipsea* (*F*_1,52_ = 5.90, *p* = 0.019; Fig. 1d), and by *Sc. stercoraria* on *Se. cynipsea* (*F*_1,36_ = 5.71, *p* = 0.022; Fig. 1f). Importantly, treatment (females only, males only, or mixed groups) was never significant for any single species pair and overall (*p* > 0.2); nor was amphibian predator identity (random effect when repeatedly used).

Figure 2 clearly demonstrates that the overall mean predator / prey size ratio (using total body length for arthropods and SVL for amphibians) is a crucial parameter influencing predation selection on prey size (cf. Brose et al., 2006): the smaller the size difference between the two interacting species, the stronger the relative viability advantage of large prey individuals, presumably primarily due to the increased handling time associated with their capture (Stephens & Krebs, 1986).

**Figure 2.**
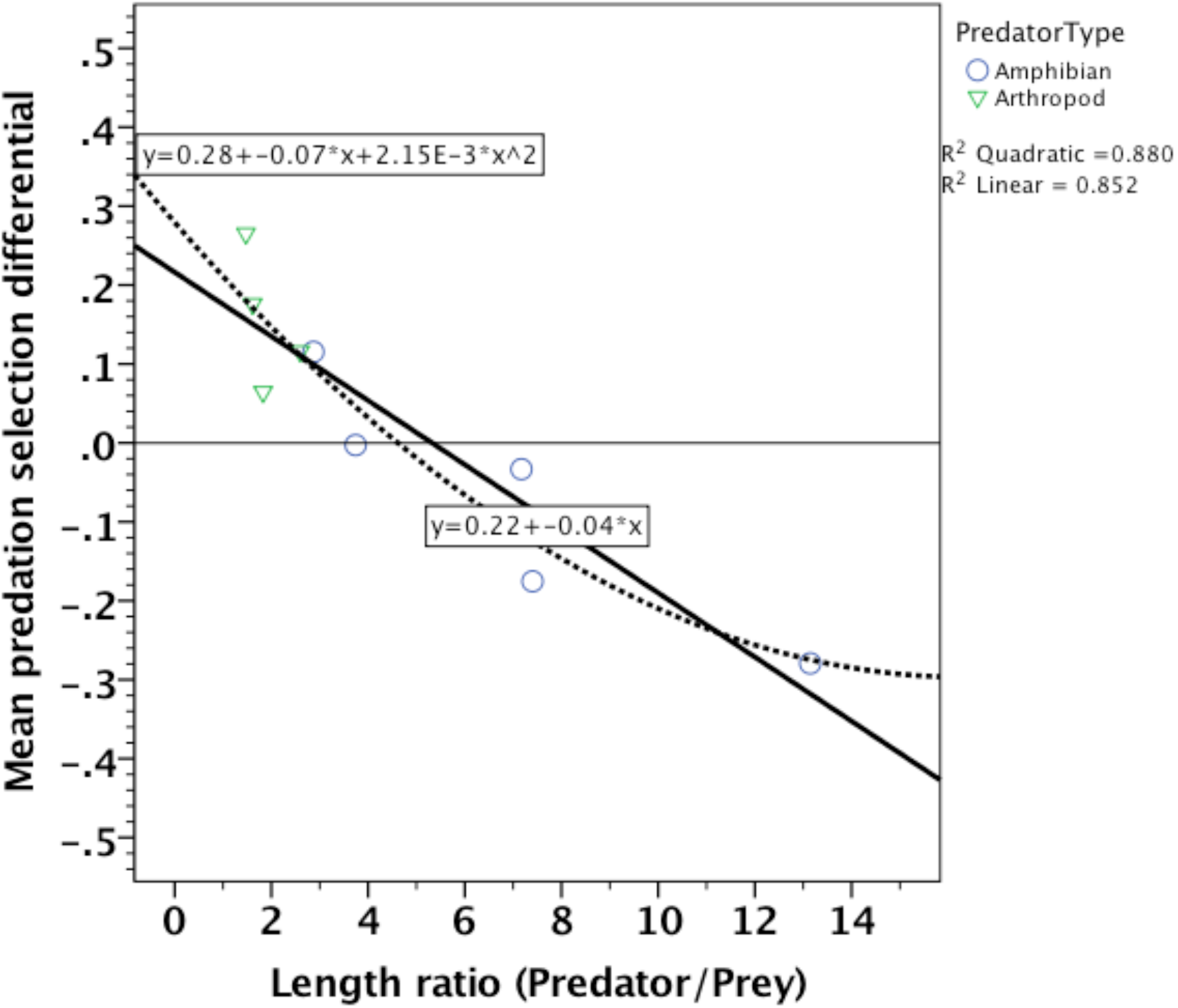
Standardized mean viability (i.e. predation) selection differential as a function of the overall average (total or SV) body length ratio between the predator and the prey. The quadratic fit is no better than the linear fit based on AIC.

Comparing the mixed-sex treatments only, we found no consistent sex bias in predation pressure by the diverse predators on the various dung fly prey species (intercept of overall model: *F*_1,113_ = 1.06, *p* = 0.308): *Rana temporaria* juveniles took significantly more male *Se. cynipsea*, while *Sc. stercoraria* took more female *Se. thoracica* (Fig. 3). The latter result was unexpected given that the orange *Se. thoracica* males were deemed much more conspicuous.

**Figure 3.**
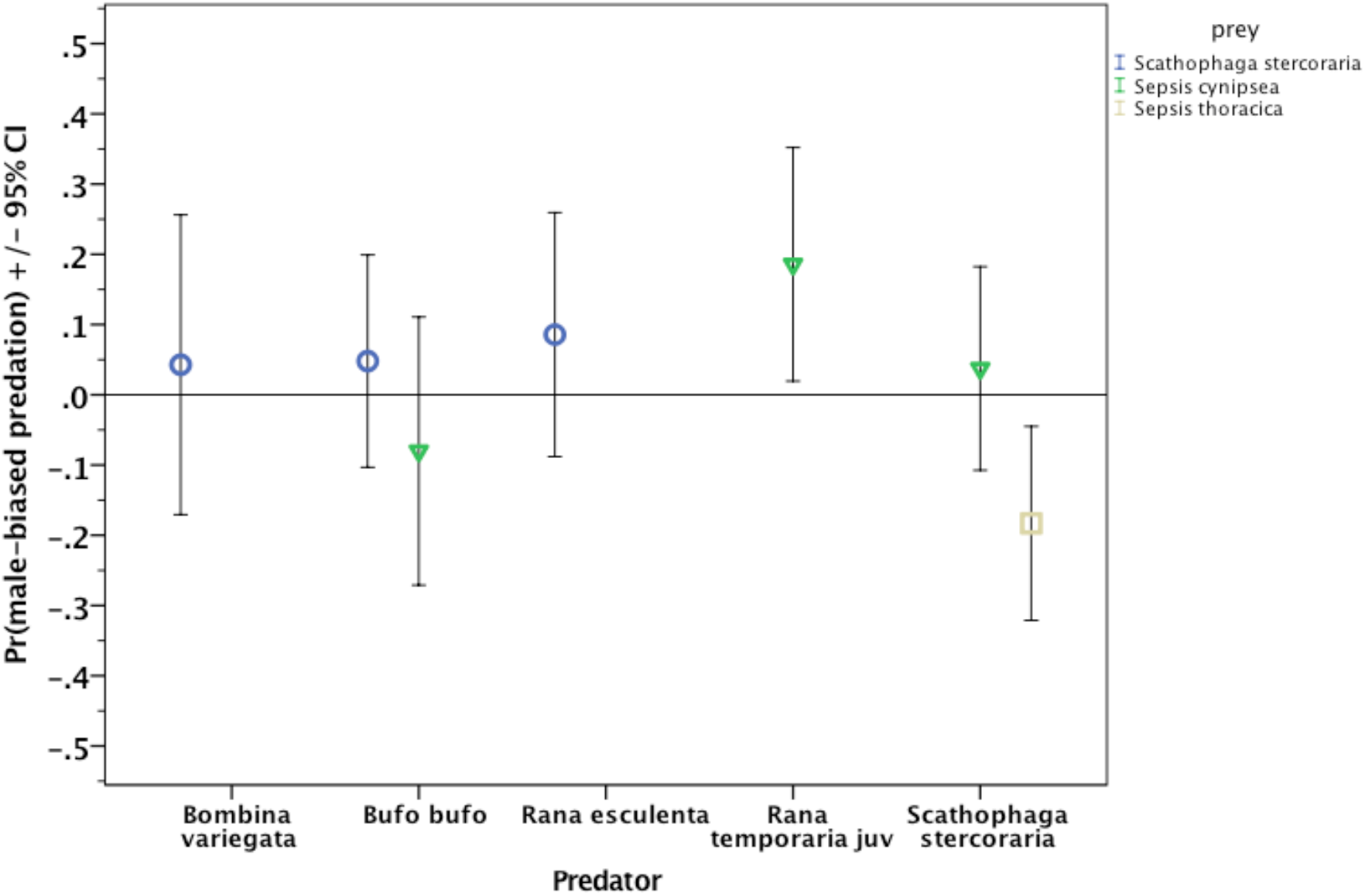
Proportional difference between the numbers of males and females eaten in the mixed-sex treatments: (m−f)/(m+f).

## DISCUSSION

Our study revealed a systematic preference of three different arthropod predators (spider, bug, fly) for smaller black scavenger flies *Se. cynipsea* and *Se. thoracica*, thus exerting positive directional viability (i.e. predation) selection on body size of the latter. In contrast, the four amphibian predators (frogs and toads) did not show a consistent prey selection pattern: the common toad *Bufo bufo* preferred to eat larger flies of two species, thus exerting negative predation selection, whereas the small common frog (*Rana temporaria*) metamorphs ate more small flies, while the two other species tested were unselective (Fig. 1). We had expected negative predation selection by the large predators (not only *Bufo bufo*), assuming optimal foraging of energetically more profitable prey with at the same time reduced handling times (Stephens & Krebs, 1986; Sih & Christensen, 2001; Brose *et al.*, 2006; Vucic-Pestic *et al.*, 2010), but this did not happen. At the same time we expected and indeed found that smaller (arthropod) predators would catch more small prey, if only because larger prey should be more effective in escaping or fending off relatively small predators, thus requiring excessive handling by the predator. Smaller individuals could have been more mobile, hence more readily detected and eaten by predators (Mühlhäuser & Blanckenhorn, 2002), but this is unlikely (and was not measured) here given generally lacking differences among the (sex) treatments (Figs. 1, 3) and lack of size effects on *Se. cynipsea* mobility in a previous study (Teuschl *et al.*, 2010). On the other hand, sepsid flies are known to release a substance smelling like geraniums suggested being a chemical deterrent against predators (Pont & Meier, 2002), of which larger individuals may release more; so this may have contributed to larger flies being eaten less.

The above-mentioned trade-off involving prey profitability and handling time here predicts opposite size-selectivity of large vs. small predators but can theoretically result in any intermediate pattern. When plotting predation selection as a function of the predator/prey length ratio (cf. Vucic-Pestic *et al.*, 2010), we indeed found a continuous shift from positive (for relatively small predators) to negative selection (for relatively large predators; Fig. 2), suggesting that the profitability aspect becomes increasingly dominant as predators increase in size. Our study generally supports similar patterns found for herbivorous insect larvae eaten by arthropods (exerting largely negatively size-dependent selection) and birds (often exerting positively size-dependent selection: Remmel *et al.*, 2011). While omitting bird predators entirely, we here focused on various common sit-and-wait amphibian and arthropod predators of widespread dung flies in grasslands, thus adding much wanted new data to the literature on size-dependent predation.

Foraging theory postulates an optimal prey size in terms of profitability (i.e. energy obtained relative to energy invested: Elner & Hughes, 1978; Stephens & Krebs, 1986), as at one point the prey becomes too big or strong for a given predator involving too much handling, or might no longer fit into the predator’s mouth (Truemper & Lauer, 2004). At the other end of the spectrum prey can also become too small for a predator, approaching invisibility, an effect that may result without necessarily invoking active selectivity as prey size falls under an assumed detectability threshold. Therefore, the relationship displayed in Fig. 2 is expected to eventually approach zero again and become curvilinear towards ever-larger predator/prey size ratios (to the right; Vucic-Pestic *et al.*, 2010), as hinted by our non-linear fit (which however is no better than the linear fit here, based on AIC). Nevertheless, at this point the prey might no longer be interesting for or even detected by many predators, ultimately resulting in no predation and hence no data, as occurred for some of our tested amphibians (cf. Table 1). Even though we initially tested several additional amphibian and arthropod predators, we perhaps necessarily ended up with a less than optimal, seemingly haphazard assortment of predator-prey pairings that behaved naturally in our experimental laboratory setting, which however should nonetheless be representative and useful in this context.

In mixed-sex groups predation risk was not higher than in single sex groups (Figs. 1, 3), generalizing similar earlier results for *Sc. stercoraria* preying upon *Se. cynipsea* (Mühlhäuser & Blanckenhorn, 2002; Teuschl *et al.*, 2010). This was unexpected, as mating is a conspicuous affair that should attract predators and at the same time distract the prey. However, while we observed some mating activity during our experiments, it was not quantified here, so we cannot address this hypothesis rigorously. We also expected predation rates to vary according to prey sex, if only because of size dimorphism, but this did not occur in general, only in the cases of *Rana temporaria* juveniles preying on *Se. cynipsea* and *Sc. stercoraria* preying on *Se. thoracica* (Fig. 3). In this context we expected the typically more mobile males to be subject to more predation than females (Teuschl *et al.*, 2010), which also did not happen: whereas *Rana temporaria* juveniles indeed took more male than female prey, this was not the case for any other predator (Fig. 3). We conclude that sex differences in predation (i.e. viability) selection by single predators appear to be rare and thus likely contribute little to sexual dimorphism (cf. Blanckenhorn, 2000, 2005).

Surprisingly, the conspicuous orange coloration of large *Se. thoracica* males did not make them more vulnerable to predation by *Sc. stercoraria*, which in fact ate more females than males, perhaps because they are more sluggish and easier to catch (Fig. 3). Although many insect and spider groups lack a receptor in the red part of the spectrum (Briscoe & Chittka, 2001; Théry & Gomez, 2010; Fabricant & Herberstein, 2015), potentially making orange-coloured prey cryptic against a green grass background, this is not generally the case for amphibians (Fite, 1976). These results suggest that, at least for the sit-and-wait predators tested here, movement seems more important than colouration (cf. Busso & Blanckenhorn, in review).

In conclusion, our study adds valuable experimental evidence to elucidating the factors influencing size-selective predation by investigating a guild of sit-and-wait amphibian and arthropod predators of insects inhabiting a common temperate grassland landscape. As each of these predators imposes a different, sex-specific selection pressure on prey body size, from the prey’s perspective viability selection in the field must be integrated across various predators to obtain net selection. This is experimentally difficult if not impossible given the multifarious predator-prey relationships in any ecosystem, many of them unspecific. Functional investigations of interactions between single predator and prey species, as conducted here, are a good start but necessarily remain incomplete. More comprehensive phenomenological studies of selection in the wild, which unfortunately typically do not consider predator species identity or investigate concrete mechanisms, thus remain indispensable when taking the prey perspective (e.g. Blanckenhorn *et al.*, 1999; Jann *et al.*, 2000).

## ACKNOWLEDGEMENTS

We acknowledge funding from the Swiss National Fund and the University of Zurich. Thanks to Daniel Hänni, Jenny Herzog, Anna Kopps, Marco Lichtsteiner and Samuel Tanner for helping with the experiments, and to Heinz-Ulrich Reyer, Christoph Vorburger, and Josh van Buskirk for providing predators.

## 3 anonymous peer reviews

### Referee: 1

Comments to the Author

This study is not particularly ambitious but it still provides valuable building blocks for our understanding of the role of predators in the evolution of (insect) body size. The study is carefully performed and the text is very well written. I have one concern about the statistical approach used (number 3) and a few suggestions with respect to presentation which the authors may wish (but need not) to consider.

#### General

1. My feeling is that the aims of the study as formulated at the end of the Introduction sound little too trivial. Perhaps for any predator, the functional relationship between prey_ size (on x-axis) and probability_to_be _predated_upon (y-axis) is bell-shaped, i.e. it approaches zero at both ends and has a maximum in between. It follows that for each predator, we can certainly get both negative and positive relationships as dependent on the range of prey sizes considered. It is quite trivial so far. Quantitative parameters of this function are not trivial, and are exciting. First, the functional relationship in your Figure 2 is very interesting. Second, I would discuss the kurtosis(?) (“flatness vs pointedness”) of the bell-shaped function. You found that, for anurans, the function is quite flat (no size dependence over a wide range of prey sizes). I would discuss this little bit. May such flatness be characteristic of a sit-and-wait predator – for a frog, there are no searching costs and the costs of handling a prey item are minimal, so it should be worth to try everything which may fit into the big mouth? Back to the beginning of what I wanted to say, I would advertise such quantitative aspects more clearly when listing the aims of the study.
2. Numerical values of such selection differentials as calculated in your study must be situation-specific. I see at least three aspects here:

a. The numerical value derived can depend on the number of flies eaten in your trial. Consider a case in which the frog always eats large flies first and you offer a mixture consisting of 4 large and 4 small flies. If the frog eats 4 flies (all large) then the size difference between the eaten and the not-eaten sample is maximal. If the fog eats either 1 or 7 flies, the difference (= calculated selection differential) must be smaller, OK?
b. The fate of an insect (eaten or not) is determined by two aspects: its detectability and its acceptability for the predator. The relative weights of detectability and acceptability as determinants of the fate certainly vary across environments, and may also be very different in your experiments and real nature.
c. I suspect that the outcome (i.e., which insects are eaten and which are not) is also dependent on whether the predators encounter the insects one at a time or has a possibility to choose between multiple ones exposed simultaneously. For these reason, I would add a cautionary note telling the reader that the numerical values of selection differentials should be treated with caution if the aim is to extrapolate those to natural conditions. The signs can - let’s hope - be extrapolated.
3. Relying on intercepts (page 12, line 12) may need attention. Intercepts may mean different things dependent on how exactly your model is parametrized. I am afraid that the most common approach (default setting in the programs) is to define a baseline level for each of the categorical independent variables (e.g. ‘female’ for the variable ‘sex’), and then to express the values for all other levels (e.g. “male”) as “baseline + specific effect”. As far as I understand, in such a case, the intercept is the value of the response variable characteristic of the observations for which all the independent variables are at their baseline values. However, it must also be possible to parametrize the model in such a way that the intercept corresponds to the grand mean of the sample. This is perhaps what you are interested in. I think that you should check (and report in the paper) that the intercept returned by your analysis is exactly what you assume it to be.

#### Minor

- I would switch two first sentences of the Abstract. Matter of taste, however.
- P 4 line 9. May “parasitoids” be too specific here? The text has not yet become insect-specific at this point.
- P 5 line 14 and elsewhere. There are two aspects of “too large prey”: 1) just failed to catch, 2) selected for to ignore.
- P 7 lines 1-2. I consider it a rule of scientific writing that the first sentence of each paragraph should be the summary of the paragraph. This is not the case here. Moreover, I am not so sure that the share of pastoral grasslands in the landscape is a good correlate of the socioeconomical developmental level of a country:-)
- P 10 line 8. This “are small” is likely thought to apply to your specific sample. If so, make this more clear, sounds too general.
- P 16 line 15-17. This sounds like an overgeneralization, please be more specific here.
- Figure 2. Text in the panels is clearly too small.

### Referee: 2

Comments to the Author

The study by Blanckenhorn et al uses laboratory experiments of amphibian and arthropod predators feeding on dung flies to explore the effects of size, sex and coloration on predation rates. While results of single predator-prey pairs are varied, on the whole there seems to be general patterns in predator-prey body size ratios. While I found the general ideas behind the experiments to be interesting, I had some reservations about the novelty of the results, and the types of experiments used to address such broad questions in ecology and evolution. Below I list some general and specific comments.

General comments:

- I found the Introduction (up to the end of page 6) quite difficult to follow, and although its tough to know exactly what is meant in a few places, I’m pretty sure I disagree with a few points made here (see my specific comments). You address some of my specific concerns at points later in the paper, but the way you make statements in the Introduction doesn’t lead the reader to think that you will quality the statements later. I also think it needs to be streamlined a lot, to make sure the background and significant of your paper is clearly laid out.
- On the whole, you haven’t convinced me that the results you present are really that novel. Fig 1 results appear to be quite idiosyncratic. Fig 2 results are interesting, but as you say it seems like you are just capturing one side of a quadratic form that is well recognized in the literature. Again with Fig 3 the results seem to be quite idiosyncratic. On the whole, it seems like what your results show is that small predators eat smaller prey than large predators. But the way you ran your experiments we don’t know at all what the mechanisms that cause these patterns are – i.e., what are the behavioral drivers that determine size-based predation?
- Related to the above point, you build up a dichotomy of predation forces driving small body size, and prey viability driving large size, but I’m not convinced. If it is true then you need to build up a more convincing argument. For example, you say repeatedly that predators will prefer large bodied prey due to energy optimization, but it could also be because of mechanical/sensory reasons.
- Its difficult to know how (and even if) these results would transfer to the field with countless other species to interact with. As you state in the manuscript, if you become too small for one predator, then you are likely going to be just the right size for the next smallest predator, and so on. In other words, how do you expect the processes to act in the full complexity of the field?
- You sum up one of my main issues with the paper in your text on page 15-16 where you say “Even though we initially tested several additional amphibian and arthropod predators, we perhaps necessarily ended up with a less than optimal, seemingly haphazard assortment of predator-prey pairings that behaved naturally in our experimental laboratory setting, which however should nonetheless be representative and useful in this context”. Working with a single species is often easy because variation is likely to be low, but obviously there are limits to the generality of the work. Using tens or more of species would give you a sufficient range in body size and taxa that general patterns should swamp the variability, However, your paper which uses just a few species is unfortunately right in the middle – where there is a lot of species-specific variation, but not sufficient species pairs to have enough statistical power to drown out the noise. I’m not sure how this could be addressed besides doing more experiments or supplementing your data with some from the literature (if it exists).
- Another issue I have is characterized by the paragraph on mixed groups in the discussion (page 16) where you are showing a size pattern, but because you didn’t study any behavior you aren’t able to draw conclusions about what mechanisms were responsible. You actually finish the manuscript by stating that other studies in the wild don’t consider concrete mechanisms, but I’m not convinced you did either.
- Lastly, while I appreciate some self citation is necessary; there is a lot in the current version and I think for very general issues/ideas you should also cite very well known seminal or otherwise important papers.

Specific comments

Page 4, Line 2: you need to define exactly what you mean by predation, as it’s a very general term.

Page 4, line 6: Of course predation features predominantly in food web and functional response theory - that’s what these theories are based on. Other examples?

Page 4, line 7-8: not sure what you mean by “predation is not easily assessed beyond the traditional and very crude food web perspective” and if I do understand you, then I disagree. The field has moved a long way from “very crude food webs”, on countless fronts.

Page 4, Line 8: I’m still not getting what the ‘major problem’ is. Highly depends on the question right? But just to imply that predation research hasn’t moved beyond crude food webs is simply not true, and ignores an enormous amount of published literature.

Page 4, Line 10: The fact that many predators are generalist is not in itself a problem - in fact this could simply things (if, for example, most trophic interactions are size-structured, and don’t depend on species identity, then that’s a simple answer).

Page 4, Line 11: I disagree. Again, the fact the predators might be generalist does not in itself mean that interactions are complex - if everything ate everything of a certain size, for example, then understanding trophic interactions would be very simple - if its not too big or too small - just eat it. What could be a simpler process?

Page 4, Line 13-14: I find this text difficult to parse - not sure exactly what you mean by underlying behavioral or ecological mechanisms. Maybe give examples?

Page 4, Line 15: what is ‘viability selection’?

Page 4, Line 18: If it really has been suggested as a major selective force, then I think cite some other papers that just the lead authors.

Page 4, Line 19: But many prey become less vulnerable to predation through ontogeny - i.e. size refugia.

Page 4, Line 21: cite some references after “… overwhelming”

Page 4, Line 22: But you just said in the sentence above that size and sex-specific predation has been suggested as a major selective force keeping animals small.

Page 4, Line 23-24: I’m not sure what you mean by this sentence, beginning with “Importantly.” Please rework.

Page 5, Line 4: But what is the first hand? From what you state here I think you are implying from the preceding sentence that eating big prey is better - but this is not clear from the way the sentence is structured.

Page 5, Line 9: It’s not clear to me yet whether you are just talking about predators, or other kinds of consumer-resource interactions as well. For example, for a parasite the selective forces are likely to be different, pushing for smaller prey size relative to predator, and thus not intermediate pre-prey size ratios. This is why I think you need to define predator earlier on.

Page 5, Line 10: what is a ‘viability advantage’?

Page 6, Line 1: You say that terrestrial studies are typically phenomenological while only citing one paper, but you cite two papers as counter exambples? Surely if something is ‘typical’ you could find more than one study?

Page 6, Line 5-9: Also, many prey are predators themselves, so evolution is not just going to act from one side.

Page 6, Line 13: What are ‘sex attracting mate’?

Page 7: most of page 7 I think could be in the materials and methods

Page 7, Line 5: I think this is the first time you talk about conspicuousness-in which case I think it should be introduced in the beginning few paragraphs, together with size and sex.

Page 8, Line 3-7: so is you expectation that small predators will eat smaller prey than larger predators? That seems obvious to me, and (as you say) will highly depend on which system you examine.

Page 8, Line 8-10: Why? More color could signify poisinous.

Page 9, Line 19-22: why didn’t these predators eat?

Page 11, line 9: how did you match individuals?

Page 12, Line 2: but hunger level can affect predation rate a lot.

Page 13, Line 3: this could be because amphibians are so much bigger than their prey, than are the arthropod predator to their prey. (i.e., amphibian predators must have a max size limit - its just you don’t get near that with the prey size you have).

Figure 1: its tough to pull out any general patterns from these panels. Is there a simple, more effective way to show these results? These just seem like raw data - hard to pull out any general l mechanism here.

### Referee: 3

Comments to the Author

Overall, I think figure is the heart of the paper. The selection gradients based on a community of foragers are really exciting. I want to call this a signal of stabilizing selection, but since there were more than one prey population in there, it’s hard to say what it adds up to for a particular prey species. Perhaps you could clarify on that point, or alter the graph to show it for each prey species? If you agree that figure 2 is the centerpiece of your story, then I also think that the paper could use a little revising through the intro in particular, as it doesn’t really draw the reader toward this centerpiece. It seems to me to get a little lost in other issues of phenomenology and size dimorphism. Perhaps if the ‘community selection’ idea is front and center, and you ask secondarily if this has something to do with choosing males or females, then it might be a bit more streamlined. I also added a few suggested edits and comments in the document. I hope you find this useful, and of course, feel free to use or ignore whatever you like.

Phenomonological predation rates (P4, L13): At this point, it is hard to know what you might mean by this. To me, rates are things per time – just measurements. There’s nothing really phenomenological or mechanistic about them. Models or theories that explain those rates might be, but I would want more explanation here about what really is wrong with the ‘rates’.

Last Intro paragraph (P8, L2): This doesn’t feel like exciting motivation. One might naturally expect a difference, so what would it mean? The motivation I am picking up on seems to be about foraging given many different predators and how those net selective forces emerge from facing diverse predator guilds?

Figure 2: This figure seems like the most important part of the ms. I think it says that when predators are large relative to their prey they tend to take the biggest prey individuals. When predators are small relative to their prey, they tend to take the smallest individuals. I think this mean that for a given prey type, the largest and smallest predators that eat the prey exert stabilizing selection on prey size. Since handling time comes after, it would be good to be clear whether you think it is that the time cost of handling slows searching or whether the predators are trying to choose prey with low handling times. Could you add a subplot to figure 2 with size ratio and say, something like fraction of prey eaten, to see if size ratio is creating a window of sorts of viable predator prey interactions? This would allow us to visually assess whether the pattern in figure 2 resides within a range of potential predation. The alternative is that the predators that didn’t forage did not because of some reason unrelated to size

P14, L2f: There are clear allometric scaling relationships between size and mobility – see Hirt et al 2017

## REFERENCES

Abrams PA, Leimar O, Nylin S, Wiklund C. 1996. The effect of flexible growth rates on optimal sizes and development times in a seasonal environment. American Naturalist 147: 381–395.

Arnold SJ, Wade MJ. 1984. On the measurement of natural and sexual selection: applications. Evolution 38: 720–734.

Berger D, Walters R, Gotthard K. 2006. What keeps insects small? Size dependent predation on two species of butterfly larvae. Evolutionary Ecology 20: 575–589.

Blanckenhorn WU. 2000. The evolution of body size: what keeps organisms small? Quarterly Review of Biology 75: 385–407.

Blanckenhorn WU. 2005. Behavioural causes and consequences of sexual size dimorphism. Ethology 111: 977–1016.

Blanckenhorn WU. 2007. Case studies of the differential equilibrium hypothesis of sexual size dimorphism in dung flies. In Sex, Size and Gender Roles. Evolutionary Studies of Sexual Size Dimorphism (ed. by D. J. Fairbairn, W. U. Blanckenhorn and T. Szekely), Oxford University Press, pp. 106–114.

Blanckenhorn WU, Morf C, Mühlhäuser C, Reusch T. 1999. Spatiotemporal variation in selection on body size in the dung fly *Sepsis cynipsea*. Journal of evolutionary Biology 12: 563–576.

Blanckenhorn WU, Mühlhäuser C, Morf C, Reusch T, Reuter M. 2000. Female choice, female reluctance to mate and sexual selection on body size in the dungfly *Sepsis cynipsea*. Ethology 106: 577–593.

Blanckenhorn WU, Hosken DJ, Martin OY, Reim C, Teuschl Y, Ward PI. 2002. The costs of copulating in the dung fly *Sepsis cynipsea*. Behavioural Ecology 13: 353–358.

Blanckenhorn WU, Fanti J, Reim C. 2007. Size-dependent energy reserves, energy utilization and longevity in the yellow dung fly. Physiological Entomology 32: 372–381.

Blanckenhorn WU, Pemberton AJ, Bussière LF, Römbke J, Floate KD. 2010. Natural history and laboratory culture of the yellow dung fly, *Scathophaga stercoraria* (L.; Diptera: Scathophagidae). Journal of Insect Science 10: 11.

Briscoe AD, Chittka L, 2001. The evolution of color vision in insects. Annual Review of Entomology 46: 471–510.

Brose U, et al. 2006. Consumer-resource body-size relationships in common food webs. Ecology 87: 2411–2417.

Brose U. 2010. Body-mass constraints on foraging behaviour determine population and food-webs. Functional Ecology 24: 28–34.

Cohen JE, Pimm SL, Yodzis P, Saldana J. 1993. Body Sizes of animal predators and animal prey in food webs. Journal of Animal Ecology 62: 67–78.

Cordts R, Partridge L. 1996. Courtship reduces longevity of male *Drosophila melanogaster*. Animal Behaviour 52: 269–278.

Elner RW, Hughes RN. 1978. Energy maximization in the diet of the shore crab, *Carcinus maenas*. Journal of Animal Ecology 47: 103–116

Fabricant SA, Herberstein ME. 2015. Hidden in plain orange: aposematic coloration is cryptic to a colorblind insect predator. Behavioural Ecology 26: 38–44.

Fite KV (ed.). 1976. The amphibian visual system: a multidisciplinary approach, pp. 203–266. New York: Academic Press.

Fraser DF, Gilliam JF. 1992. Nonlethal impacts of predator invasion: facultative suppression of growth and reproduction. Ecology 73: 959–970.

Hammer O. 1941. Biological and ecological investigations on flies associated with pasturing cattle and their excrement. Videnskabelige Meddelelser fra Dansk Naturhistorisk Forening 105: 140–393.

Hedrick AV, Dill LM. 1993. Mate choice by female crickets is influenced by predation risk. Animal Behaviour 46: 193–196.

Jann P, Blanckenhorn WU, Ward PI. 2000. Temporal and microspatial variation in the intensities of natural and sexual selection in the yellow dung fly *Scathophaga stercoraria*. Journal of evolutionary Biology 13: 927–938

Jormalainen V, Merilaita S, Riihimäki J. 2001. Costs of intersexual conflict in the isopod *Idotea baltica*. Journal of evolutionary Biology 14: 763–772.

Kalinkat G, Rall BC, Vucic-Pestic O, Brose U. 2011. The allometry of prey preferences. Plos One 6: e25937.

Kalinkat G, Schneider FD, Digel C, Guill C, Rall BC, Brose U. 2013. Body masses, functional responses and predator-prey stability. Ecology Letters 16: 1126–1134.

Kozlowski J. 1992. Optimal allocation of resources to growth and reproduction: implications for age and size at maturity. Trends in Ecology & Evolution 7: 15–19.

Kotiaho J, Alatalo RV, Mappes J, Parri S, Rivero A. 1998. Male mating success and risk of predation in a wolf spider: a balance between sexual and natural selection? Journal of Animal Ecology 67: 287–291.

Leimar O. 1996. Life history plasticity: Influence of photoperiod on growth and development in the common blue butterfly. Oikos 76: 228–234.

Lima SL, Dill LM. 1990. Behavioural decisions made under the risk of predation: a review and prospectus. Canadian Journal of Zoology 68: 619–640.

Lüning J. 1992. Phenotypic plasticity of *Daphnia pulex* in the presence of invertebrate predators morphological and life history responses. Oecologia 92: 383–390.

Magnhagen C. 1991. Predation risk as a cost of reproduction. Trends in Ecology & Evolution 6: 183–185.

Mänd T, Tammaru T, Mappes J. 2007. Size-dependent predation risk in cryptic and conspicuous insects. Evolutionary Ecology 21: 485–498.

Mühlhäuser C, Blankenhorn WU. 2002. The costs of avoiding matings in the dung fly *Sepsis cynipsea*. Behavioural Ecology 13: 359–365.

Parker GA. 1970. The reproductive behaviour and the nature of sexual selection in Scathophaga stercoraria L. (Diptera: Schathophagidae) I. Diurnal and seasonal changes in population density around the site of mating and oviposition. Journal of Animal Ecology 39: 185–204.

Parker GA. 1972. Reproductive behaviour of *Sepsis cynipsea* (L.) (Diptera - Sepsidae). II. The significance of the precopulatory passive phase and emigration. Behaviour 41: 242–250.

Partridge L, Farquhar M. 1981. Sexual activity reduces lifespan of male fruitflies. Nature 294: 580–582.

Pont AC, Meier R. 2002. The Sepsidae (Diptera) of Europe. Fauna Entomologica Scandinavica 37: 1–221.

Pomiankowski A. 1987. The costs of choice in sexual selection. Journal of Theoretical Biology 128: 195–218.

Reim C, Teuschl Y, Blanckenhorn WU. 2006. Size-dependent effects of larval and adult food availability on reproductive energy allocation in the yellow dung fly. Functional Ecology 20: 1012–1021.

Remmel T, Davison J, Tammaru T. 2011. Quantifying predation on folivorous insect larvae: the perspective of life-history evolution. Biological Journal of the Linnean Society 104: 1–18.

Remmel T, Tammaru T. 2009. Size-dependent predation risk in tree-feeding insects with different colouration strategies: A field experiment. Journal of Animal Ecology 78: 973–980.

Rice JA, Crowder LB, Rose KA. 1993. Interactions between size-structured predator and prey populations: experimental test and model comparison. Transactions of the American Fisheries Society 122: 481–491.

Roff DA. 1992. The evolution of life histories, theory and analysis. London (UK): Chapman and Hall.

Rowe L. 1994. The cost of mating and mate choice in waterstriders. Animal Behaviour 48: 1049–1056.

Ruxton GD, Sherratt TN, Speed MP. 2004. Avoiding Attack: The Evolutionary Ecology of Crypsis, Warning Signals and Mimicry. Oxford: Oxford University Press.

Sandrock C, Schirrmeister BE, Vorburger C. 2011. Evolution of reproductive mode variation and host associations in a sexual-asexual complex of aphid parasitoids. BMC Evolutionary Biology 11: 348.

Scherer AE, Smee DL. 2016. A review of predator diet effects on prey defensive responses. Chemoecology 26: 83–100.

Sih A, Christensen B. 2001. Optimal diet theory: when does it work and when and why does it fail? Animal Behaviour 61: 379–390.

Skidmore P. 1991. Insects of the British Cow-Dung Community. Slough, United Kingdom: Richmond Publishing Co.

Slagsvold T, Lifjeld JT, Stenmark G, Breiehagen T. 1988. On the cost of searching for a mate in female pied flycatchers *Ficedula hypoleuca*. Animal Behaviour 36: 433–442.

Sogard SM. 1997. Size-selective mortality in the juvenile stage of teleost fishes: A review. Bulletin of Marine Science 60: 1129–1157.

Stearns SC, Koella J. 1986. The evolution of phenotypic plasticity in life history traits: predictions of reaction norms for age and size at maturity. Evolution 40: 893–914

Stephens DW, Krebs JR. 1986. Foraging Theory. Princeton (NJ): Princeton University Press.

Stireman JO, Singer MS. 2003. Determinants of parasitoid-host associations: insights from a natural tachinid-lepidopteran community. Ecology 84: 296–310.

Svanbäck R, Eklöv P. 2011. Catch me if you can—predation affects divergence in a polyphenic species. Evolution 65: 3515–3526.

Symondson WOC, Sunderland KD, Greenstone MH. 2002. Can generalist predators be effective biocontrol agents? Annual Review of Entomology 47: 561–594.

Teuschl Y, Reim C, Blanckenhorn WU. 2010. No size-dependent reproductive costs in black scavenger fly (*Sepsis cynipsea*) males. Behavioural Ecology 21: 85–90.

Théry M, Gomez D, 2010. Insect colours and visual appearance in the eyes of their predators. Advances in Insect Physiology 38: 267–353.

Truemper HA, Lauer TE. 2004. Gape limitation and piscine prey size-selection by yellow perch in the extreme southern area of Lake Michigan, with emphasis on two exotic prey items. Journal of Fish Biology 66: 135–149.

Vucic-Pestic O, Rall BC, Kalinkat G, Brose U. 2010. Allometric functional response model: Body masses constrain interaction strengths. Journal of Animal Ecology 79: 249–256.

Ward PI. 1983. The effects of size on the mating behaviour of the dung fly *Sepsis cynipsea*. Behavioural Ecology & Sociobiology 13: 75–80.

Ward PI, Hemmi J, Röösli T. 1992. Sexual conflict in the dung fly *Sepsis cynipsea*. Functional Ecology 6: 649–653.

Warncke E, Terndrup U, Michelsen V, Erhard A, 1993. Flower visitors to Saxifraga hirculus in Switzerland and Denmark, a comparative study. Botanica Helvetica 103: 141–147.

Watson PJ, Arnqvist G, Stallmann RR. 1998. Sexual conflict and the energetic costs of mating and mate choice in water striders. American Naturalist 151: 46–58.

Wellborn GA. 1994. Size-biaised predation and prey life histories: a comparative study of freshwater amphipod populations. Ecology 75: 2104–2117.

Werner EE, Anholt BR. 1993. Ecological consequences of the trade-off between growth and mortality mediated by foraging activity. American Naturalist 142: 242–272.

Zuk M, Kolluru GR. 1998. Exploitation of sexual signals by predators and parasitoids. Quarterly Review of Biology 73: 415–438.

